# 5-hydroxymethylcytosine profiling of cell-free DNA identifies bivalent genes that are prognostic of survival in high-risk neuroblastoma

**DOI:** 10.1101/2023.04.27.538309

**Authors:** Mohansrinivas Chennakesavalu, Kelley Moore, Gepoliano Chaves, Sahil Veeravalli, Rachel TerHaar, Tong Wu, Ruitu Lyu, Alexandre Chlenski, Chuan He, Andrea Piunti, Mark A. Applebaum

**Author notes:** Corresponding Author: Mark A. Applebaum, MD Department of Pediatrics University of Chicago 5841 S. Maryland Ave., MC4060 Chicago, IL 60637 Facsimile.

## Abstract

Neuroblastoma is the most common extra-cranial solid tumor in childhood and epigenetic dysregulation is a key driver of this embryonal disease. In cell-free DNA from neuroblastoma patients with high-risk disease, we found increased 5-hydroxymethylcytosine (5-hmC) deposition on Polycomb Repressive Complex 2 (PRC2) target genes, a finding previously described in the context of bivalent genes. As bivalent genes, defined as genes bearing both activating (H3K4me3) and repressive (H3K27me3) chromatin modifications, have been shown to play an important role in development and cancer, we investigated the potential role of bivalent genes in maintaining a de-differentiated state in neuroblastoma and their potential use as a biomarker. We identified 313 genes that bore bivalent chromatin marks, were enriched for mediators of neuronal differentiation, and were transcriptionally repressed across a panel of heterogenous neuroblastoma cell lines. Through gene set variance analysis, we developed a clinically implementable bivalent signature. In three distinct clinical cohorts, low bivalent signature was significantly and independently associated with worse clinical outcome in high-risk neuroblastoma patients. Thus, low expression of bivalent genes is a biomarker of ultra-high-risk disease and may represent a therapeutic opportunity in neuroblastoma.

## INTRODUCTION

Neuroblastoma is an embryonal malignancy of neural crest origin and is the most common extra-cranial solid tumor in childhood ^1^. Neuroblastoma is notable for its biological heterogeneity, manifesting in a broad spectrum of clinical behavior. Reflecting this, low-risk neuroblastoma accounts for one of the highest rates of spontaneous regression of all human cancers, yet those with high-risk (HR) disease have a long-term survival rate of approximately 40-50% ^2^. Improved biomarkers are necessary to identify patients who are unlikely to respond to current therapy regimens, allow for more precise treatment plans, and identify patients who should be considered for alternative approaches ^3^.

Unlike most cancers of adults, neuroblastoma is characterized by a low mutational burden, and epigenetic dysregulation is a key driver of oncogenesis ^4–6^. Most DNA methylation occurs on cytosines in the context of Cytidine-phosphate-Guanosine (CpG) dinucleotides ^7^. 5-hydroxymethylcytosine (5-hmC) is a stable intermediate generated in the process of removing methyl groups from cytosines and accumulates in enhancers, gene bodies, and promoters ^8, 9^. This conversion of 5-methylcytosine (5-mC) to 5-hmC, catalyzed by TET proteins, is associated with open chromatin and active gene expression in multiple cancers, including neuroblastoma ^10, 11^. With the introduction of nano-hmC-seal technology, genome-wide 5-hmC profiles can be generated in a cost effective and sensitive manner ^12^. We previously showed that 5-hmC profiles from diagnostic neuroblastoma tumors are prognostic of outcome ^11^. Subsequently, we showed that circulating cell-free DNA (cfDNA) derived 5-hmC profiles identified patients with neuroblastoma who experienced subsequent relapse with high sensitivity and specificity ^13^.

Based on these studies, 5-hmC profiles from both neuroblastoma tumors and cfDNA offer prognostic value. However, the relationship between 5-hmC profiles from tumor biopsies and cfDNA is yet to be elucidated in neuroblastoma. cfDNA molecules are released into plasma through apoptosis, necrosis, and active secretion and reflect the genome and epigenome of their cells of origin ^14^. Tumor cells with the highest metastatic potential are thought to contribute disproportionately to cfDNA in plasma ^15^. Because the half-life of cfDNA is on the order of hours, cfDNA is dynamic and may highlight the most fundamental mechanisms of aggressive tumor biology. Thus, we hypothesized that differential 5-hmC modifications selectively enriched in cfDNA compared to diagnostic tumor biopsy samples from patients with neuroblastoma would identify epigenetic changes associated with aggressive tumor behavior and help identify novel biomarkers for identifying high-risk neuroblastoma patients who may benefit from alternate therapeutic approaches.

## RESULTS

### Genome-wide 5-hmC profiles reveal higher levels of 5-hmC on PRC2 targets in patient derived cfDNA compared to tumor DNA

To identify epigenetic drivers of aggressive neuroblastoma behavior, we compared genome-wide 5-hmC profiles from neuroblastoma patient derived cfDNA and tumor biopsy samples (dbGaP accession number: phs001831.v1.p1). Principal Component Analysis (PCA) revealed that principal component 1 (PC1), correlating to sample type (cfDNA or tumor), explained 97% of the variation among cfDNA and tumor 5-hmC profiles (**SUPPLEMENTAL FIGURE 1A)**. Using the Estimate of Stromal and Immune cells in Malignant Tumors using Expression data (ESTIMATE) algorithm, relative cellular heterogeneity among cfDNA and tumor biopsy samples was quantified^16^. As expected, cfDNA samples had significantly higher Immune Scores than tumor biopsy samples (p < 2.2e^-16^), while tumor biopsy samples had significantly higher Stromal Scores than cfDNA samples (p = 2.8e^-14^) (**SUPPLEMENTAL FIGURE 1B,C**) ^16^. Incorporating PC1, Immune Score, and Stromal Score as covariates, we identified 7,792 genes with significantly different 5-hmC deposition (false discovery rate (FDR) < 0.05) between cfDNA from HR patients with active disease (n=64) and HR tumors from diagnostic biopsies (n=48) (**FIGURE 1A**). Using genes with log-fold-change in the top 10% of differentially hydroxymethylated genes (n=1,559 genes) between cfDNA and tumor samples, unsupervised clustering of 129 cfDNA samples clearly differentiated samples from patients with the highest metastatic burden from samples derived from patients with lower/no metastatic burden, suggesting that the identified genes correlated with aggressive neuroblastoma phenotype (**SUPPLEMENTAL FIGURE 1D**).

**FIGURE 1:**
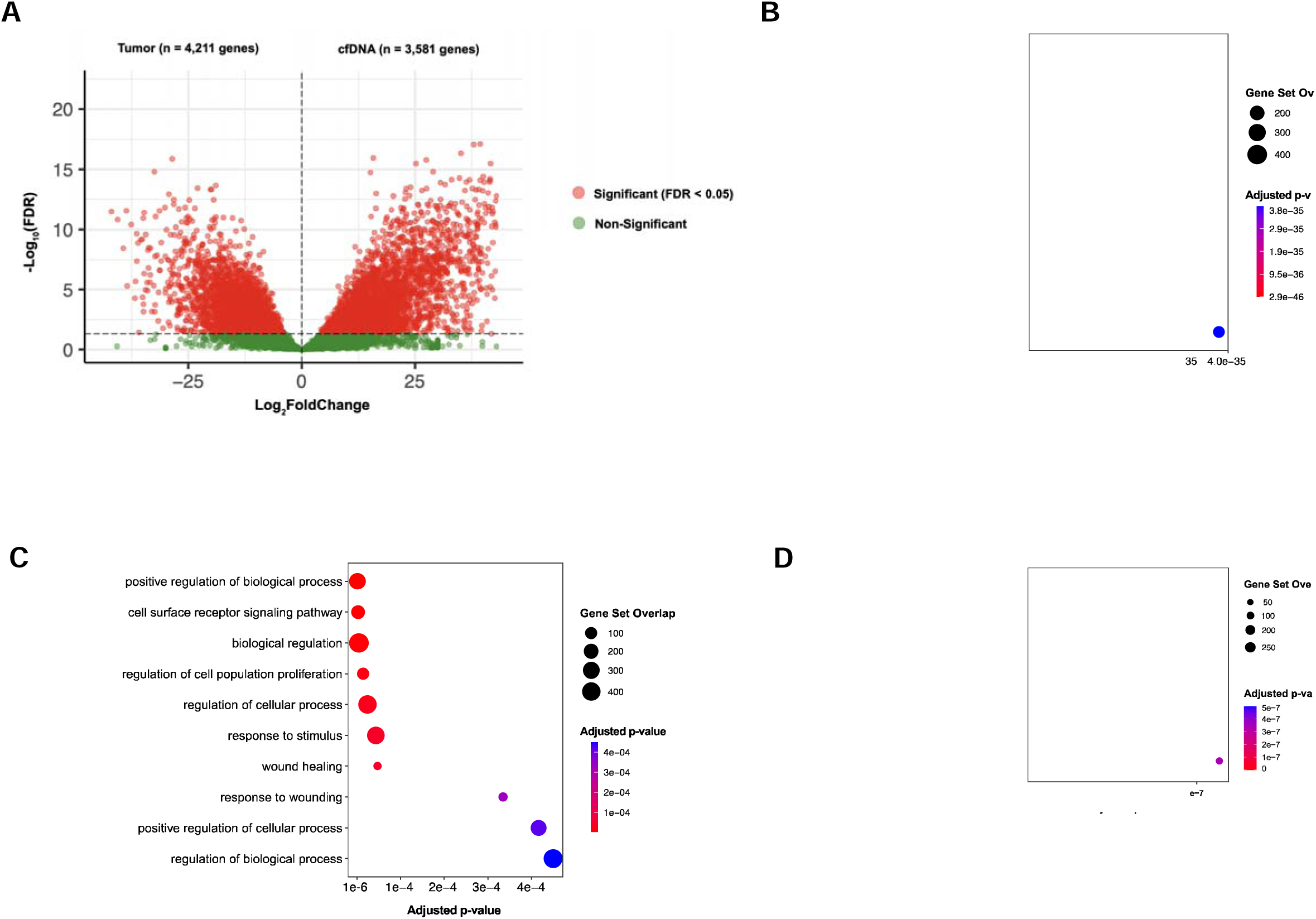
Comparison of cfDNA and tumor 5-hmC profiles highlights increased 5-hmC on PRC2 target genes in neuroblastoma. Differential hydroxymethylated genes across cfDNA and tumor samples, as visualized through **A)** Volcano plot. 3,581 genes had significantly greater 5-hmC deposition in cfDNA samples, while 4,211 genes had significantly greater 5-hmC in tumor samples. **B)** Gene set enrichment analysis of top 10 biological pathways significantly (FDR < 0.05) enriched in cfDNA 5-hmC profiles and **C)** tumor 5-hmC profiles. Gene set enrichment analysis demonstrates significant (FDR < 0.05) enrichment of 5-hmC on **D)** REST, SUZ12, and EZH2 target genes in cfDNA.

Next, we performed gene set enrichment analysis to identify biological pathways differentially hydroxymethylated between cfDNA and tumor DNA samples ^17, 18^. Genes in the top 10% by log-fold-change with significantly increased 5-hmC deposition in cfDNA compared to tumor DNA (n=780) were enriched for Gene Ontology Biological Processes including nervous system development, neurogenesis, and neuron differentiation, pathways highly relevant to neuroblastoma (**FIGURE 1B**). Conversely, genes in the top 10% by log-fold-change with significantly increased 5-hmC deposition in tumor DNA compared to cfDNA were enriched for pathways including cell surface receptor signaling pathway and wound healing (**FIGURE 1C)**.

We also observed significant enrichment of 5-hmC on SUZ12, EZH2, and REST target genes in cfDNA (**FIGURE 1D**). SUZ12 and EZH2 are both subunits of Polycomb Repressive Complex 2 (PRC2), while REST is a transcription factor that has previously been shown to interact with PRC2 ^19, 20^. PRC2 is an enzymatic complex that mediates the mono-, di-, and trimethylation of lysine 27 of histone 3 (H3K27me1-2-3), and H3K27me3 is a histone posttranslational modification associated with compacted chromatin and transcriptional silencing ^21^. PRC2, through its catalytic activity, is involved in cell differentiation, identity, and stem-cell plasticity, and has been shown to play a role in neuroblastoma pathogenesis ^20, 22^. Prior studies have shown that 5-hmC is associated with PRC2 on bivalent genes, a class of genes involved in numerous developmental and oncogenic contexts characterized by the presence of both repressive (H3K27me3) and activating (H3K4me3) chromatin marks ^23–27^. Given our findings of increased 5-hmC on PRC2 targets in aggressive neuroblastoma, we next sought to define and characterize the biological relevance of bivalent genes in neuroblastoma.

### Bivalent genes are enriched for mediators of differentiation and are repressed in neuroblastoma cell lines

To determine the biologic ramifications of the bivalent chromatin state in neuroblastoma, we established the deposition of a panel of chromatin marks across two neuroblastoma cell lines: SK-N-BE2 and NBLW-N. MACS2 was used to define H3K27me3 and H3K4me3 peaks within 5kb of the transcription start site (TSS) of protein-coding genes in these cell lines ^28^. In SK-N-BE2, we identified 1,672 genes that were enriched for H3K27me3. Of the genes enriched for H3K27me3, 1,148 genes were enriched for H3K27me3-only and 524 genes enriched for both H3K27me3 and H3K4me3, a histone modification found at the promoter of actively transcribed genes, which were defined as bivalent ^26^. Similarly, in NBLW-N, we identified 2,830 genes that were enriched for H3K27me3. Of the genes with H3K27me3 deposition, 1,901 genes were enriched for H3K27me3-only and 928 genes were bivalent. There was significant overlap between the sets of genes identified in SK-N-BE2 and NBLW-N, with 313 bivalent genes (p < 0.0001) and 806 H3K27me3-only genes (p < 0.0001) shared between the two cell lines.

To further characterize the distribution of bivalent genes and H3K27me3-only genes, we investigated the deposition of 5-hmC and H3K27ac, generally deposited in euchromatin regions ^20^, and single stranded DNA (ssDNA) using KAS-Seq, across the two gene sets in relation to highly expressed genes ^29, 30^. Highly expressed genes were defined independently in each included cell line as the 2,000 genes with the highest transcripts per million (TPM). In the NBLW-N cell line, both H3K27me3-only and bivalent genes had little to no presence of markers of open chromatin (H3K27ac deposition) and single stranded DNA (ssDNA) compared to highly expressed genes, confirming low chromatin accessibility (**FIGURE 2A-D**). Both H3K27me3-only and bivalent genes had deposition of 5-hmC at the TSS, while highly expressed genes had a notable lack of 5-hmC at the TSS. Bivalent genes had significantly higher deposition of 5-hmC at the TSS compared to H3K27me3-only genes (p = 0.002; **FIGURE 2E**), consistent with prior studies conducted in embryonic stem cells (ESCs) ^24^. Similar results were observed in the SK-N-BE2 cell line (**SUPPLEMENTAL FIGURE 2**). Genes lacking H3K27me3 had the highest expression, as expected (**FIGURE 2F**). Though both bivalent and H3K27me3-only genes were relatively transcriptionally repressed, bivalent genes had significantly higher expression across a panel of eight cell lines compared to those with H3K27me3-only (**FIGURE 2F**).

**FIGURE 2:**
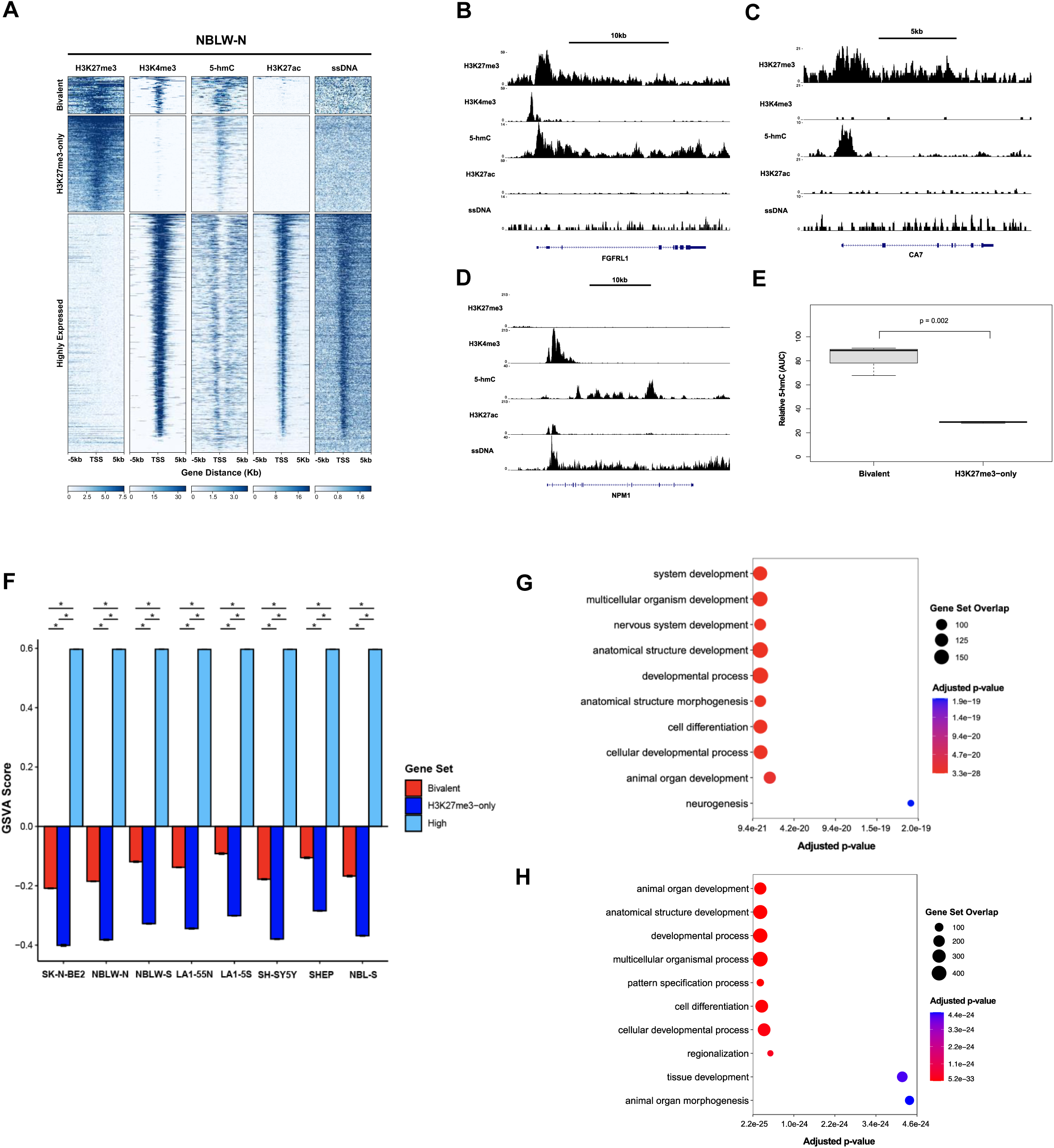
Bivalent genes are enriched for 5-hmC at the TSS, transcriptionally repressed, and enriched for mediators of neuronal differentiation. **A)** H3K27me3, H3K4me3, 5-hmC, H3K27ac, and ssDNA profiles (normalized to RPKM) +/-5kb of the TSS, in the NBL-W-N cell line. Profiles were obtained across three different gene lists: bivalent (313 genes), H3K27me3-only genes (806 genes), and highly expressed genes (2000 genes) in the NBLW-N cell line. H3K27me3, H3K4me3, 5-hmC, H3K27ac, and ssDNA tracks are shown for representative **B)** bivalent (FGFRL1), **C)** H3K27me3-only (CA7), and **D)** highly expressed (NPM1) genes in the NBLW-N cell line. H3K27me3, H3K4me3, and H3K27ac are scaled to one another, while ssDNA and 5-hmC are separately scaled to one another. **E)** Relative 5-hmC within 5kb of the TSS, as measured by area under the curve (AUC), is shown for bivalent genes and H3K27me3-only genes in the NBLW-N cell line. Significance was determined by two-tailed t-test. **F)** Gene set variance analysis (GSVA) scores of bivalent genes, H3K27me3-only genes, and highly expressed genes. Significance determined by two-tailed t-test. Error bars denote mean +/-one standard deviation. * denotes p-value < 0.05. **G)** Biological pathways enriched in bivalent genes, as determined by GSEA. **H)** Biological pathways enriched in H3K27me3-only genes, as determined by GSEA.

Next, we investigated the biologic significance of bivalent genes in neuroblastoma ^26^. Through gene set enrichment analysis, we found that both H3K27me3-only and bivalent genes were similarly significantly enriched for biologic pathways involved in early embryonal development, including pathways such as “animal organ development,” “developmental process,” and “cellular developmental process” (**FIGURE 2G,H**) ^17^. Notably, “nervous system development” and “neurogenesis”, pathways aberrantly repressed in HR neuroblastoma tumors, were among the most significant biological pathways enriched in bivalent genes and were not present in the top ten most significant biological pathways enriched in H3K27me3-only genes. This suggests that repression of bivalent genes may be involved in preventing the terminal differentiation of neuroblastoma cells into neurons, a critical mechanism involved in the clinical response of HR neuroblastoma tumors to therapy ^31^.

### Bivalent gene expression predicts survival in high-risk neuroblastoma

Given the biological relevance of bivalent genes, we investigated the clinical significance of these genes in three distinct clinical cohorts of neuroblastoma ^32–34^. Utilizing gene set variance analysis (GSVA), bivalent signature scores were determined for each included sample ^35^. In the SEQC-NB dataset, which includes 498 neuroblastoma patient samples, high-risk (HR) tumors (n=176) had significantly lower bivalent signature scores (p < 2.2e-16) compared to non-high-risk (non-HR) tumors (n=322) (**FIGURE 3A**) ^32^.

**FIGURE 3:**
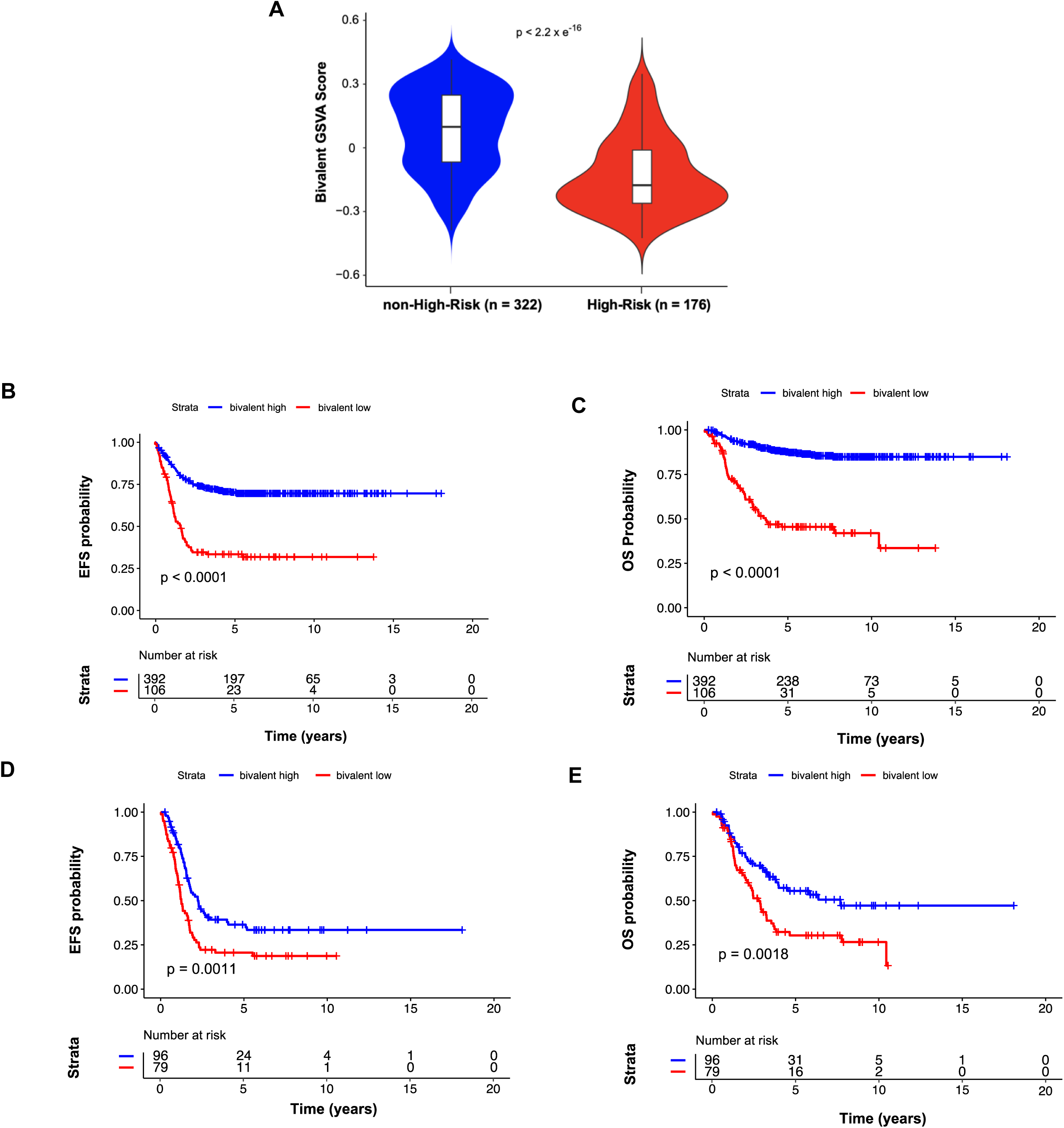
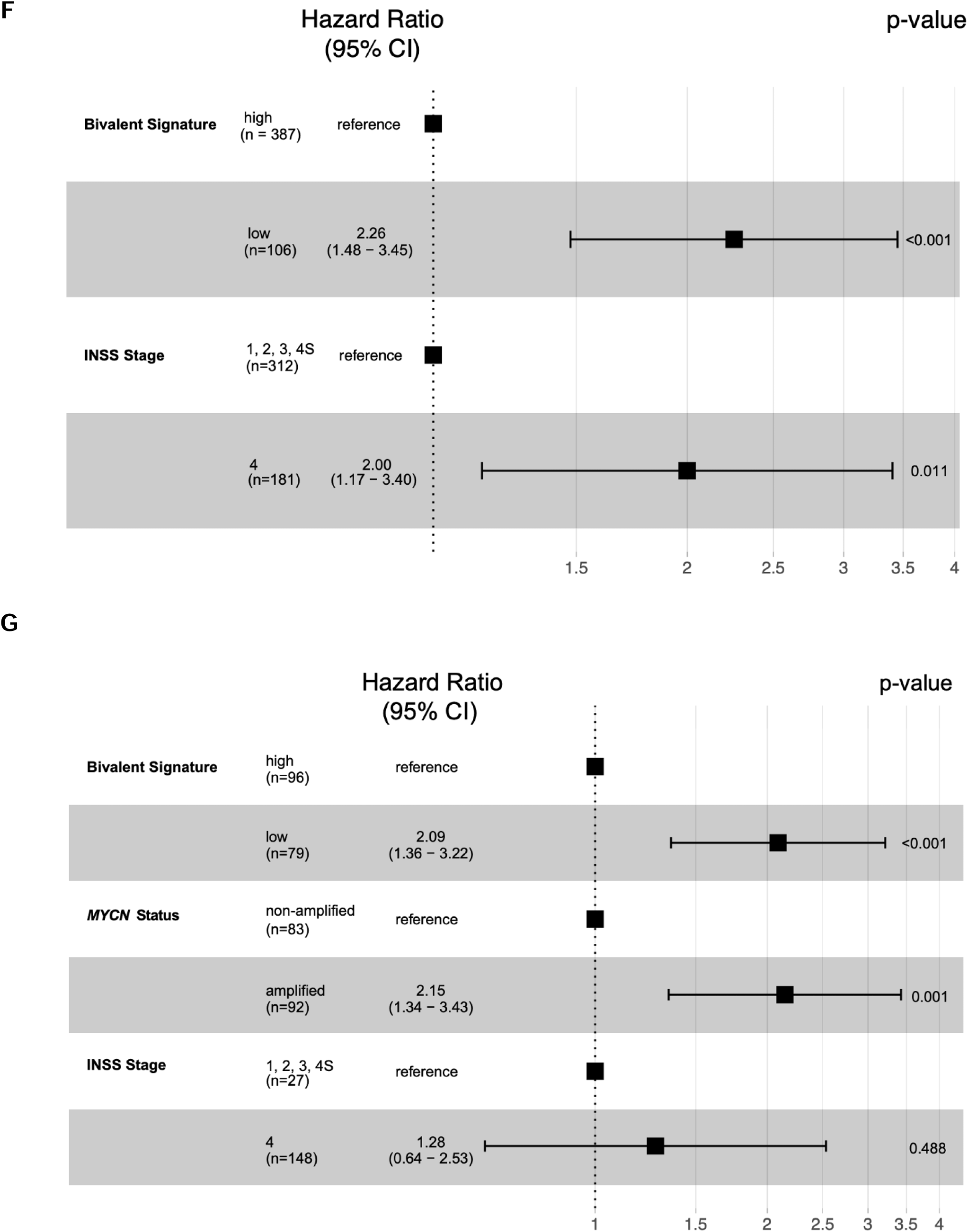
Low expression of bivalent genes is associated with poor clinical outcome in neuroblastoma patients. Expression and phenotypic data for 498 neuroblastoma patients (SEQC-NB cohort) were obtained from R2. **A)** Bivalent GSVA scores were compared across non-High-Risk (n=322 tumors) and High-Risk (HR) (n=176 tumors). Two-tailed t-test was used to determine statistical significance. Kaplan-Meier curves depicting **B)** event-free (EFS) and **C)** overall survival (OS) in all 498 patients by relative bivalent gene expression. Kaplan-Meier curves depicting **D)** EFS and **E)** OS in HR tumors with known *MYCN-*amplification status (n=175) by relative bivalent gene expression. Samples with relative enrichment of bivalent genes are labeled as “bivalent high”, while samples with relative repression of bivalent genes are labeled as “bivalent low”. Significance for Kaplan-Meier curves determined by log-rank test. **F)** Multivariable Cox regression analysis for OS in 493 patients with known clinical and biological variables, stratified by *MYCN*-amplification status and age at diagnosis. **G)** Multivariable Cox regression analysis for OS in 175 HR patients with known clinical and biological variables, stratified by age at diagnosis.

The ability of bivalent signatures scores to predict overall survival (OS) was evaluated through receiver operating characteristic (ROC) curve analysis. In the full SEQC-NB cohort, the area under the curve (AUC) of bivalent signatures for OS was 75.6% [95% CI: 70.3% - 80.8%] (**SSUPPLEMENTAL FIGURE 3A**). In the cohort of SEQC-NB HR tumors, AUC of bivalent signatures for OS was 60.1% [95% CI: 51.7% - 68.6%] (**SUPPLEMENTAL FIGURE 3B**). To assess the potential of bivalent signatures to identify patients with worse outcome, the optimal bivalent signature cutoff score (- 0.195) was determined through interrogation of the ROC curve of SEQC-NB HR tumors. Using this cutoff point, tumors were categorized as “bivalent-low” or “bivalent-high.” In the full SEQC-NB cohort, tumors with low bivalent signatures had significantly worse event-free survival (EFS) (5-year EFS: 33.4% [95% CI: 25.3%-44.1%] vs 70.3% [95% CI: 65.8%-75.1%]; p < 0.0001) and OS (5-year OS: 45.5% [95% CI: 36.3%-56.9%] vs 87.3% [95% CI: 83.9%-90.8%]; p < 0.0001) compared to tumors with high bivalent signature scores (**FIGURE 3B,C**). In the subset of HR patients in the SEQC-NB cohort, tumors with low bivalent signature scores also had significantly worse EFS (5-year EFS: 20.6% [95% CI: 13.1%-32.4%] vs 36.4% [95% CI: 27.5%-48.2%]; p = 0.0011) and OS (5-year OS: 30.3% [95% CI: 21.0%-43.8%] vs 55.5% [95% CI: 45.5%-67.7%]; p = 0.0018) compared to tumors with high bivalent signature scores (**FIGURE 3D,E**).

Low bivalent signature was significantly associated with worse OS in univariate Cox regression analysis in both the full SEQC-NB (hazard ratio: 5.78 [95% CI: 3.92- 8.51], p < 2.2e-16) and HR SEQC-NB (hazard ratio: 1.92 [1.27-2.91], p = 0.00215) cohorts. Multivariable Cox regression analysis including standard clinical measures of aggressive disease (*MYCN* amplification status, risk status, age at diagnosis, International Neuroblastoma Staging System [INSS] stage) revealed that a low bivalent signature score was independently associated with worse OS in both the full SEQC-NB (hazard ratio: 2.26 [95% CI: 1.48-3.45], p = 0.000167) and HR SEQC-NB (hazard ratio: 2.09 [1.36-3.22], p = 0.000815) cohorts (**FIGURE 3F,G**).

Similar results were observed in an independent dataset consisting of 105 neuroblastoma tumor samples obtained at diagnosis (GSE73518) ^33^. Bivalent signatures predicted OS with high accuracy in both the full GSE73518 cohort (n=105) and the HR GSE7518 (n=56) cohort with AUC’s of 72.2% [95% CI: 61.4% - 83.0%] and 79.1% [95% CI: 67.3% - 90.9%], respectively (**SUPPLEMENTAL FIGURE 3C,D**). Using the same bivalent signature cutoff score determined in the HR SEQC-NB cohort, we found that tumors with low bivalent signature scores had significantly worse EFS (5-year EFS: 26.2% [95% CI: 12.9%-52.9%] vs 58.9% [95% CI: 48.9%-71.0%]; p = 0.00028) and OS (5-year OS: 43.0% [95% CI: 26.6%-69.6%] vs 79.8% [95% CI: 71.1%-89.6%]; p < 0.0001) compared to tumors with high bivalent signature scores in the full GSE73518 cohort (**FIGURE 4A,B**). Furthermore, within the HR subset of the GSE73518 cohort, tumors with low bivalent signature scores also had significantly worse EFS (5-year EFS: 12.5% [95% CI: 3.4%-45.7%] vs 41.5% [95% CI: 28.3%-60.9%]; p = 0.00017) and OS (5-year OS: 12.5% [95% CI: 2.5%-63.5%] vs 59.2% [95% CI: 44.4%-78.8%]; p < 0.0001) compared to tumors with high bivalent signature scores (**FIGURE 4C,D**). Multivariable Cox regression analysis confirmed that low bivalent signature score was significantly independently associated with worse OS in both the full GSE73518 (hazard ratio: 4.93 [95% CI: 2.21-11.01], p = 0.00010) and HR GSE73518 (hazard ratio: 4.77 [95% CI: 2.11-10.76], p = 0.000167) cohorts (**FIGURE 4E,F**).

**FIGURE 4:**
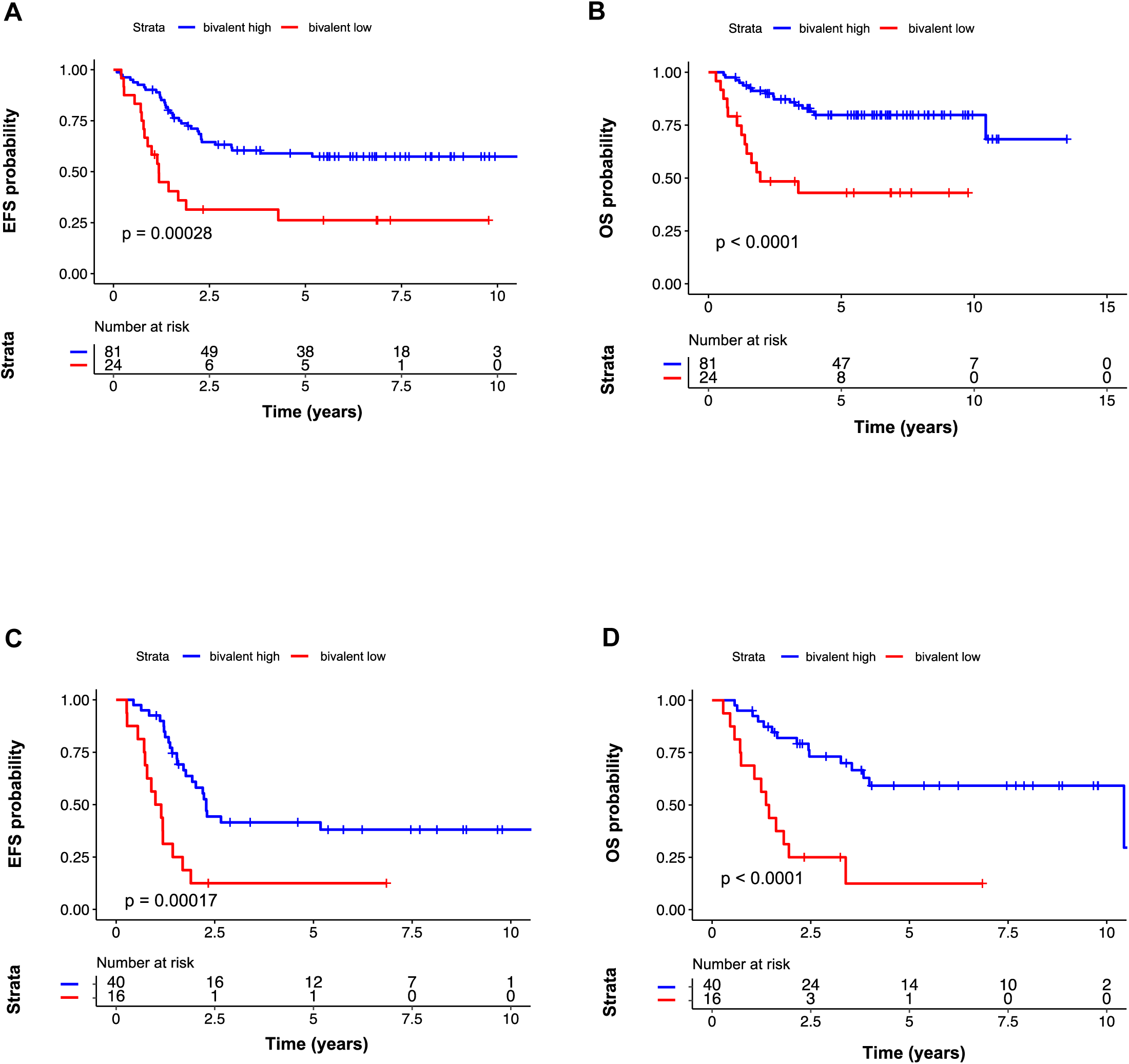
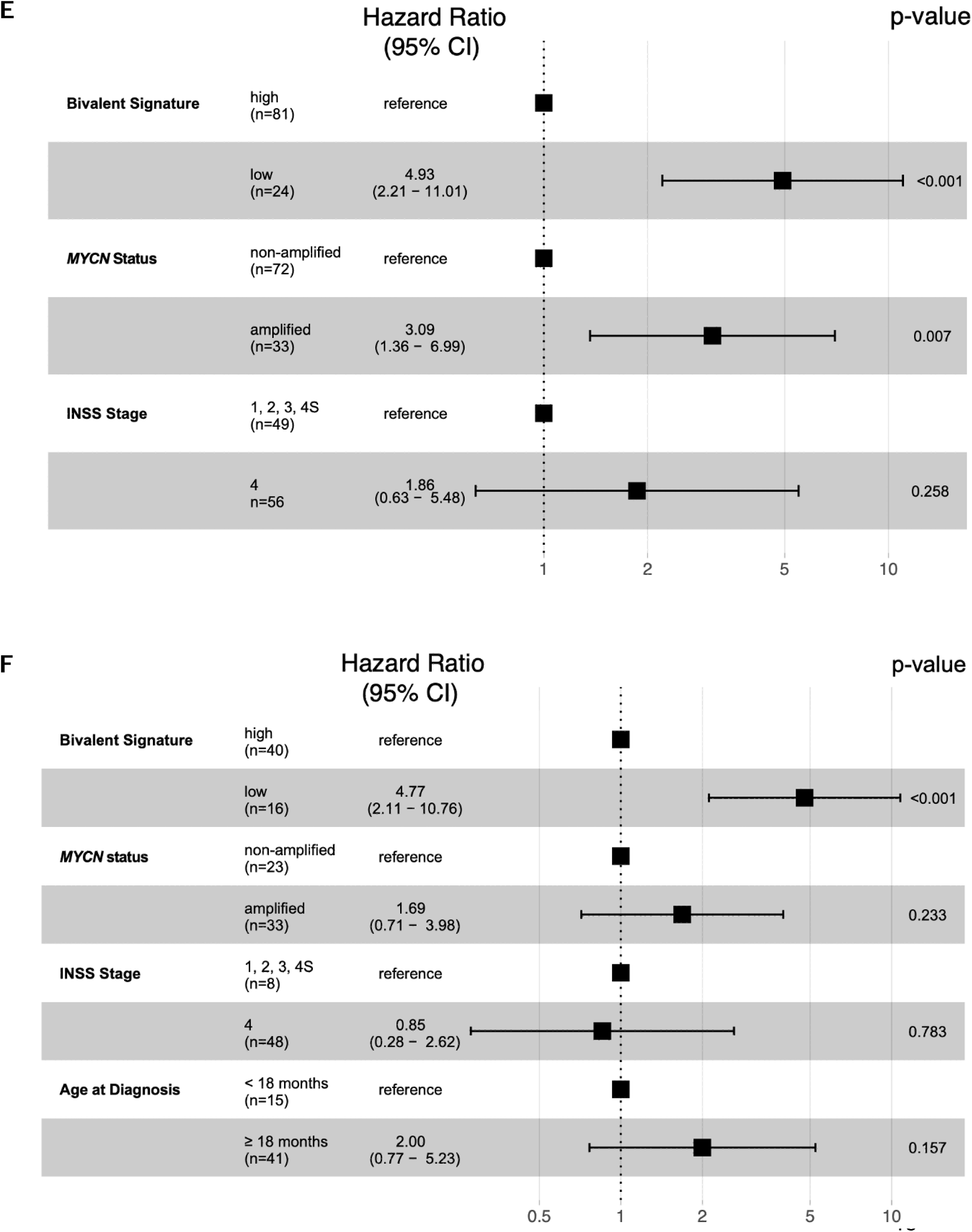
Low expression of bivalent genes is associated with poor clinical outcome in neuroblastoma patients in the GSE73517 dataset. Expression and phenotypic data for 105 neuroblastoma patients (GSE73517) were obtained from R2. Kaplan-Meier curves depicting **A)** event-free (EFS) and **B)** overall survival (OS) in all 105 patients by relative bivalent gene expression. Kaplan-Meier curves depicting **C)** EFS and **D)** OS in HR tumors (n=56) by relative bivalent gene expression. Samples with relative enrichment of bivalent genes are labeled as “bivalent high”, while samples with relative repression of bivalent genes are labeled as “bivalent low”. Significance for Kaplan-Meier curves determined by log-rank test. **E)** Multivariable Cox regression analysis for OS in all 105 patients, stratified by age at diagnosis. **F)** Multivariable Cox regression analysis for OS in 56 HR patients.

Finally, in a third distinct cohort of neuroblastoma patients, using the same cutoff established in the HR subset of the SEQC-NB cohort, we observed that low expression of bivalent genes was associated with worse OS in the E-MTAB-8248 (n=130) and the HR subset of the E-MTAB-8248 cohorts (n=41) (**SUPPLEMENTAL FIGURE 4A,B)** ^34^.

Following multivariable Cox regression analysis, low bivalent signature was again independently associated with worse OS in both the E-MTAB-8248 cohort (hazard ratio: 3.95 [95% CI: 1.28-12.17], p = 0.0165) and HR subset of the E-MTAB-8248 cohorts (hazard ratio: 27.80 [95% CI: 2.67-289.6], p = 0.000193) (**SUPPLEMENTAL FIGURE 4C,D**).

## DISCUSSION

In this study, we found increased 5-hmC accumulation on PRC2 target genes in cfDNA compared to diagnostic tumor biopsy samples from patients with neuroblastoma. We also show significant enrichment of 5hmC at 313 bivalent genes that were transcriptionally repressed and enriched for mediators of neuronal differentiation.

Through GSVA, we developed a clinically implementable bivalent signature. In three distinct clinical cohorts, HR patients with low bivalent signature scores had significantly worse survival compared to those with high bivalent signature scores, highlighting the robust nature of this gene signature. Finally, multivariable Cox modeling demonstrated low expression of bivalent genes was associated with worse OS independent of validated clinical and biological prognostic factors. Improved understanding of the epigenomic underpinnings of HR neuroblastoma is of crucial importance for optimal risk stratification and the development of targeted therapies, and the use of bivalent gene signatures may enable further refinements in therapeutic decision making.

A novel approach to identify epigenomic drivers of HR neuroblastoma was used in our study through analysis of patient cfDNA derived 5-hmC profiles. cfDNA profiling represents a uniquely powerful tool to study the biology of neuroblastoma. In comparison to other pediatric solid tumors, patients with neuroblastoma have significantly higher amounts of cfDNA circulating in plasma ^36^. Additionally, our prior work and that of others have demonstrated that tumor-specific alterations can be monitored during the course of therapy through cfDNA surveillance and that cfDNA profiling captures dynamic tumor evolution in a way not possible with standard tumor biopsies ^13, 37, 38^. Importantly, in the current study we demonstrated that a comparison of cfDNA and diagnostic tumor derived 5-hmC profiles identified a set of differentially hydroxymethylated genes associated with worse clinical outcome, supporting the hypothesis that cfDNA samples are enriched for DNA from the most aggressive, rapidly dividing tumor cells. The identification of this biology from cfDNA highlights the potential of cfDNA 5-hmC profiling to uncover fundamental biological mechanisms in neuroblastoma and pediatric solid tumors more generally.

We found that 5-hmC deposition at PRC2 targets in cfDNA was associated with aggressive clinical features in neuroblastoma. While 5-hmC deposition is generally correlated with open chromatin and active gene expression, previous studies demonstrated a dual role for TET1 and 5-hmC in mouse ESCs. In mouse ESCs, 5-hmC is enriched at the gene body of transcriptionally active genes and also at the promoter region in PRC2 repressed developmental regulators ^39^. Furthermore, TET1 directly interacted with PRC2 in ESCs, but not more differentiated cells ^40^. In this context, our findings raise the possibility of a similar dual role for 5-hmC in HR neuroblastoma as mechanisms utilized in ESCs to maintain a dedifferentiated, stem-like state may also be utilized in HR neuroblastoma to promote malignancy and prevent tumors from differentiating into neurons. Further mechanistic studies are required to clarify the direct relationship between 5-hmC and PRC2 and its potential oncogenic role in neuroblastoma. Additional understanding of this relationship may pave the way for new targeted therapies to block the potential interaction between 5-hmC and PRC2 to enable neuronal differentiation.

Although numerous recent studies have characterized the open chromatin and enhancer landscape in neuroblastoma, less is known about the repressive landscape, and to date, the bivalent gene landscape of neuroblastoma has yet to be explored ^41–43^. Bivalent genes are defined by the presence of histone modifications associated with both activation (H3K4me3) and repression (H3K27me3) at the promoter region, and are generally transcriptionally inactive or transcribed at very low levels ^26, 27^. Bivalent genes were initially discovered in the context of developmentally regulated genes in ESCs, and were posited to be “poised” for rapid activation during development ^23^. Though direct evidence validating this model remains elusive, bivalency has been suggested to be important in embryogenesis, differentiation, and the maintenance of genome architecture ^44^. Bivalent domains in glioblastoma have been shown to identify a subtype-independent signature of early neural development, while bivalent domains in ovarian cancer were prone to hypermethylation at relapse after chemotherapy ^45, 46^. In the current study, we found that bivalent genes were enriched for mediators of neuronal differentiation and were transcriptionally repressed. Though we identified bivalent genes using two *MYCN*-amplified, adrenergic neuroblastoma cell lines, low expression of bivalent genes was conserved across a panel of eight heterogenous neuroblastoma cell lines, including *MYCN*-non-amplified and mesenchymal type cell lines, suggesting that the bivalent genes we identified are not lineage-specific. This is further supported by the finding that bivalency was still prognostic in patients after accounting for *MYCN*-amplification, a marker associated with more adrenergic tumors ^47^. Thus, aberrant silencing of bivalent genes may represent a previously unexplored mechanism by which a de-differentiated state is maintained in neuroblastoma.

The current Children’s Oncology Group risk classification for patients with neuroblastoma utilizes a combination of clinical and biological features including stage, age at diagnosis, *MYCN*-amplification status, tumor ploidy, tumor histology, and presence or absence of segmental chromosomal aberrations ^48^. Currently, no clinical or biologic features are utilized to further stratify patients with HR disease. Here, we showed that low expression of bivalent genes was significantly associated with worse OS in three distinct cohorts of HR patients, independent of known risk factors. Our results suggest a robust association between bivalent gene expression and survival in neuroblastoma, wherein low expression of this gene set could be used as a clinical biomarker to identify patients with ultra-high-risk disease who are unlikely to benefit from standard cytotoxic therapy and may be prioritized for novel approaches.

Methodologic limitations exist in our approach to defining bivalency using independent H3K4me3 and H3K27me3 chromatin immunoprecipitation followed by sequencing (ChIP-Seq) experiments. Through this approach, we are unable to directly quantify amounts of H3K27me3 in relation to H3K4me3, nor are we able to determine whether these marks co-localize at a sub-cellular or mono-nucleosomal level.

Furthermore, we identified bivalent genes in neuroblastoma cell lines rather than patient derived tumor biopsies. Through the use of reICeChIP, a calibrated sequential ChIP method, a recent study has called into question the bivalent hypothesis, suggesting that bivalency is widespread at many thousands of promoters, does not resolve with differentiation, and are not transcriptionally poised ^44^. However, reassuringly, we find that the identified bivalent gene set is more highly expressed than genes only bearing H3K27me3, as assessed through bulk RNA-sequencing, and robustly predicts survival in neuroblastoma patients and is thus likely biologically relevant in this disease. In future studies, we will utilize reICeChIP on DNA derived from diagnostic tumor biopsies to re-examine the presence of bivalent genes in neuroblastoma.

In summary, our results contribute to a growing understanding of the role of epigenetic dysregulation in the development of neuroblastoma. We used insights from patient derived cfDNA 5-hmC profiles to identify bivalent genes as mediators of neuronal differentiation in neuroblastoma, and we found that low expression of bivalent genes was associated with worse survival across three distinct cohorts of neuroblastoma patients. As these bivalent genes are potential regulators of a stem-like state, we highlight a novel, unexplored mechanism by which dedifferentiated states may be maintained in HR neuroblastoma that may have clinical implications as a prognostic biomarker.

## METHODS

### Tumor biopsy derived genome-wide 5-hmC profile analysis

Raw 5-hmC sequencing data and phenotypic information for n=129 cfDNA and n=107 tumor samples were obtained from dbGaP accession number: phs001831.v1.p1. Raw read quality was assessed using fastqc (version 0.11.5) with default settings ^49^. Trimmomatic (version 0.36) was used to filter raw reads with a minimum Phred score of 33 ^50^. Reads were aligned to GRCh38 using Bowtie2 (version 2.3.0) with default settings ^51^. Duplicate reads were removed using picard (version 2.18.29) with default settings ^52^. Finally, aligned reads were counted across an entire gene body using featureCounts of subread (version 1.5.3), with the gene flag ^53^.

To account for cellular heterogeneity in comparing cfDNA and tumor biopsy samples, the Estimate of Stromal and Immune cells in Malignant Tumors using Expression data (ESTIMATE) package (version 1.0.13) was utilized in R (version 3.6.3) to generate Immune Scores, Stromal Scores, and ESTIMATE scores for all samples ^16^. Principal component analysis (PCA) was conducted using ggfortify (version 0.4.14) ^54^.

To identify genes with differential levels of 5-hmC modifications across samples, 5-hmC count data were loaded into the DESeq2 (version 1.28.1) package in R (version 4.2.0) ^55^. Differential 5-hmC analysis was performed with sex, batch, Principal Component 1, Immune Score, and Stromal Score as covariates in the model. An FDR of less than 0.05 was considered significant. Significantly differentially 5-hydroxymethylated genes were visualized with EnhancedVolcano (version 1.10.0) ^56^. Unsupervised clustering analysis was performed using the ComplexHeatMap package (version 2.4.3) in R ^57^. ENRICHR, g:Profiler, and clusterProfiler were utilized to conduct gene set enrichment analysis of significant genes with the 10% greatest fold change between groups ^17, 18, 58^.

### Cell Culture

Neuroblastoma cell lines LA1-55n, LA1-5s, SH-SY5Y, SHEP, NBL-W-N, NBL-W-S, SK-N-BE2, and NBL-S were grown at 5% CO_2_ in RPMI1640 medium (Life Technologies) supplemented with 10% heat-inactivated FBS, 2 mmol/L l-glutamine, and 1% penicillin/streptomycin. LA1-55n, LA1-5s, SH-SY5Y, SHEP, NBL-W-N, NBL-W-S, SK-N-BE2, and NBL-S were kind gifts from Dr. Susan Cohn (University of Chicago). Cells were verified with STR testing and were confirmed to be free of mycoplasma. For ChIP-seq, cells were counted using a hemocytometer and grown in a T225 flask for a period of 24 hours. Cells were harvested from the flask, pelleted in 50ml conical tubes, washed with 1x DPBS solution (Gibco 14190-144) and resuspended in 20 ml RPMI 1640 medium (Life Technologies). Resuspended cells were separated into 10ml aliquots for chromatin extraction. For hmC-seal, cells were counted using a hemocytometer and grown in a two T75 flasks to a confluency of 50%. Then one T75 flask for each cell line was grown at 5% CO_2_ for 48 hours. 9 × 10^6^ cells were harvested from each flask, pelleted in 50 ml conical tubes, washed with 1x DPBS solution (Gibco 14190-144) and resuspended in Qiagen Cell Lysis Solution (Qiagen).

### Chromatin Extraction and ChIP-seq

Cells were fixed with 1% formaldehyde (Sigma F87750) for 10 minutes at room temperature and quenched with 0.125M glycine (Boston BioProducts C4375). Cells were pelleted, resuspended in an equal volume of 1x PBS (Gibco 10010-023), and snap frozen in liquid nitrogen. Cell pellets were thawed, lysed with 5mL lysis buffer 1 (50mM HEPES-KOH pH 7.5, 140 mM NaCl, 1mM EDTA, 10% Glycerol, 0.5% NP-40, 0.25% Triton-X), and mixed at 4°C for 10 minutes. Cells were pelleted, lysed with 5mL lysis buffer 2 (200mM NaCl, 1 mM EDTA, 0.5 mM EGTA, 10mM Tris-HCl pH8) and mixed at 4°C for 10 minutes. Cells were pelleted and lysed with 500µl lysis buffer 3 (1mM EDTA, 0.5mM EGTA, 10mM Tris-HCl pH8, 100mM NaCl, 0.1% Na-Deoxycholate, 0.5% N-lauroyl sarcosine, 1% Triton). Lysates were sonicated using a Diagenode Bioruptor (30 seconds on and 30 seconds off) for 10 minutes in precooled conditions. Lysates were pelleted at 4°C and the supernatant was collected. A 5% aliquot of extracted chromatin was used as an input control and incubated at 65°C overnight for reverse crosslinking. 250µl of lysis buffer 3 was added to the remaining lysate which was then snap frozen. The input control was purified using Qiagen PCR kit (Qiagen 28104) and 1µg of extracted chromatin was run on a 2% agarose gel to visualize 100-400bp sheared DNA.

For immunoprecipitation, solubilized chromatin was thawed on ice and pelleted at 13,000g for 5 minutes. The supernatant was incubated with either 5µg of Anti-H3K27me3 antibody (Abcam ab6002), 5µg of Anti-H3K4me3 antibody (Cell Signaling Technologies C42D8), or 5µg Anti-Histone H3K27ac antibody (Abcam ab4729) and rotated overnight at 4°C. Dynabeads™ Protein G (Invitrogen 10004D) were used to capture chromatin fragments. Dynabeads™ were furthered washed with lysis buffer 3, lysis buffer 3 with 1M NaCl, lysis buffer3 with 0.5M NaCl, and TE (10mM Tris pH 8, 1mM EDTA). DNA was eluted from beads using elution buffer (1% SDS, 250 mM NaCl, 10mM Tris-HCl pH8, 1mM EDTA) at 65°C for 1 hour, treated with RNAase A (Qiagen 158922) at room temperature for 1 hour and Proteinase K (Qiagen 158918) at 55°C for 2 hours followed by a reverse cross-link at 65°C overnight. DNA was purified using Qiagen PCR kit (Qiagen 28104). ChIP DNA libraries were made using Ovation® Ultralow System V2 kit (Tecan Genomics 0344NB-32) and sequenced on an Illumina NovaSeq 6000 in single-end 80-bp mode.

### hmC-Seal Protocol

Genomic DNA from cells in Qiagen Cell Lysis were purified using Qiagen Puregene Cell Kit (Qiagen 158722) according to manufacturer’s instructions and resuspended in 100µl nuclease-free H_2_0. 10µg of purified gDNA was diluted to a final volume of 130µl in nuclease-free H_2_0. gDNA was sonicated twice using a Diagenode Bioruptor (30 seconds on and 30 seconds off) on high setting for 10 minutes in precooled conditions to an average size of 100-400 bp. The sonicated volume of gDNA was then adjusted to a volume of 20µl using a SpeedVac Vacuum Concentrator. Glucosylation reactions were performed in a 50µl solution containing sonicated DNA with UDP-Azide-Glucose (Active Motif 55020), T4 beta-glucosyltransferase and 10X EPi Buffer (Fisher Scientific FEREO0831) and incubated at 37°C for 2 hours. Reactions were purified using Micro Bio-Spin™ P-6 Gel Columns, Tris Buffer (Bio Rad 7326221). Columns were washed 4X with 500ul H_2_O before purification. The modified DNA was added to 1µl [1.0mM] Disulfide Biotin DBCO (Click Chemistry Tools A112-5) and incubated at 37°C for 2 hours. The DNA was purified using Qiagen QiaQuick Kit (Qiagen 28104) according to manufacturer’s instructions and eluted in 50µl of Qiagen Elution Buffer. Invitrogen™ Dynabeads™ MyOne™ Streptavidin C1 beads (Fisher Scientific 65-001) were washed 3X with 1X B&W buffer diluted from 2X B&W buffer (10mM Tris HCl pH 7.5, 1mM EDTA, 2M NaCl, 0.01% Tween 20). Beads were resuspended in 50µl 2X B&W buffer, added to purified DNA, and incubated for 1 hour with top-over-end rotation. Beads were collected on a magnetic stand and washed 3X with 1X B&W buffer. Beads were then resuspended in 70µl [100mM] DTT (Thermo Scientific R0861) for 1 hour at 37C with top-over-end rotation. Beads were collected on a magnetic stand and the hmc-enriched DNA in the supernatant was collected and purified using Micro Bio-Spin™ P-6 Gel Columns, Tris Buffer (Bio Rad 7326221). DNA was purified again using Qiagen MinElute Kit (Qiagen 28004) according to the manufacturer’s instructions and eluted in 12µl of nuclease-free H_2_0. DNA libraries were made using Ovation® Ultralow System V2 kit (Tecan Genomics 0344NB-32) according to manufacturer’s instructions and sequenced on the Illumina NextSeq 500 instrument.

### KAS-seq library construction

KAS-seq libraries of purified genomic DNA were generated as described ^29, 30^. Briefly, cells were incubated for 5–10 min at 37 °C, 5% CO2 in complete culture medium containing 5 mM N3-kethoxal. Using the PureLink genomic DNA mini kit (Thermo, K182002), genomic DNA (gDNA) was isolated from collected cells. 1ug gDNA was suspended in 95uL DNA elution buffer with 5uL 20 mM DBCO-PEG4-biotin (DMSO solution, Sigma, 760749), 25 mM K3BO3 while being gently shaken at 37°C for 1.5 hours. 5µL RNase A (Thermo, 12091039) was added to the reaction mixture, which was then incubated at 37°C for 5 minutes. The DNA Clean & Concentrator-5 kit (Zymo, D4013) was used to recover biotinylated gDNA. gDNA was suspended in 100µL water and fragmented to 150-350 bp size using the Bioruptor Pico (30s-on/30s-off) for 30 cycles. 5% of fragmented gDNA was used as input, while the remaining 95% was used to enrich biotin-tagged DNA through incubation at room temperature with Dynabeads MyOne Streptavidin C1 (Thermo, 65001). The beads were washed, and DNA was eluted by heating the beads at 95°C for 10 minutes in 15µL H_2_O. Accel-NGS Methyl-seq DNA library kit (Swift, 30024) was used for library construction on eluted DNA and corresponding input. The Illumina Nextseq500 platform was used to sequence libraries with single-end 80-bp mode.

### RNA sequencing and RNA library construction

RNA was isolated using TRIzol® reagent (Life Technologies) according to the manufacture’s protocol. The concentration was measured using UV spectroscopy (DeNovix). Extracted RNA was sent to the Genomics Facility at the University of Chicago for library preparation and sequencing. RNA quality and quantity was assessed using the Agilent bio-analyzer. Strand-specific RNA-SEQ libraries were prepared using an oligo-dT based RNA-SEQ library protocol (Illumina/NEB/Kapa provided). Library quality and quantity was assessed using the Agilent bio-analyzer and libraries were sequenced using an Illumina NovaSeq 6000.

### Sequencing data processing, analysis, and visualization

Raw read quality for all sequencing data was data was assessed using fastqc (version 0.11.5) with default settings ^49^. For cell line ChIP-Seq and 5-hmC data, Trimmomatic (version 0.36) was used to filter raw reads with a minimum Phred score of 33 ^50^. Reads were aligned to GRCh38 using Bowtie2 (version 2.3.0) with default settings ^51^. Duplicate reads were eliminated using picard (version 2.18.29) with default settings ^52^. MACS2 (version 2.1.0) with standard settings was utilized to define H3K4me3 enriched regions, while the “--broad” flag was used to define H3K27me3 enriched regions ^28^. Relative ChIP occupancy signal for H3K27me3 was defined using deepTools “bamCompare,” and H3K27me3 enriched genes were defined as genes with greater than a 2 fold increase in ChIP signal compared to the input sample ^59^. Relative ChIP occupancy signal for H3K4me3 was defined using deepTools “bamCompare,” and H3K4me3 enriched genes were defined as genes with increased ChIP signal compared to the input sample. bigwig files normalized to reads per kilobase million (RPKM) were generated using deepTools (version 2.0), and tracks were visualized using UCSC Genome Browser ^59, 60^. Genome-wide ChIP-Seq and 5-hmC profiles were visualized using ngs.plot (version 2.63) and deepTools (version 2.0) ^59, 61^. Significance of overlap between gene sets were determined through hypergeometric t-test.

For cell line RNA-Seq data, Trimmomatic (version 0.36) was used to process raw reads, and reads were aligned to GRCh38 using STAR-aligner (version 2.6.1d) ^50, 62^. Aligned reads were counted using featureCounts of subread (version 1.5.3), with the exon flag.^53^ Raw counts were converted to transcripts per million (TPM) for downstream analysis. Highly expressed genes for each of the eight included cell lines were defined as the top 2000 genes with the highest TPM. GSVA scores for bivalent genes, H3K27me3-only genes, and highly expressed genes within each cell line were determined and compared via two-tailed t-tests.

For cell line KAS-seq raw data, reads were trimmed using the trim-galore package in single-end mode ^63^. Trimmed reads were aligned to hg38 using bowtie2 (v2.3.0) with standard parameters and sorted to bam files using samtools sort (v1.9) ^51, 64^. samtools rmdup (v1.9) was utilized to obtain unique mapped reads, which were then extended to 150 bp. Bam files were visualized with deepTools ^59^.

### Gene expression and survival analysis in neuroblastoma tumors and cell lines

Expression data and associated phenotypic data for 498 neuroblastoma tumors (SEQC-NB cohort) were downloaded from R2 (r2.amc.nl) and are also available at GSE49710 and GSE62564 ^32, 65^. Similarly, expression data and associated phenotypic data for a separate dataset of 105 neuroblastoma tumors were downloaded from R2 and are also available at GSE73517 ^33^. Finally, expression data and associated phenotypic for a third data set of 223 neuroblastoma tumors was downloaded from R2 and are also available at E-MTAB-8248 ^34^. Due to concern for repeated samples between the SEQC-NB and E-MTAB-8248 cohorts, samples in the E-MTAB-8248 with the same age at diagnosis, MYCN-amplification status, and INSS Stage as those found in the SEQC-NB cohort were excluded, resulting in a remaining dataset of 130 tumors. Because risk classification was not available in the E-MTAB-8248 cohort, samples from patients with INSS Stage 4 disease and an age at diagnosis of over 18 months were defined as HR (n=41 samples).

GSVA was performed in R using the *GSVA* package (version 1.46.0) on tumor samples using our bivalent gene list ^35^. AUROC analyses examining the predictive power of bivalent GSVA scores for OS were implemented in R using the pROC package (version 1.18.0) ^66^. Optimal bivalent GSVA score cutoff (-0.195) was determined in the HR-only subset of the SEQC-NB cohort (n=176 tumors) through the “thresholds = “best”” parameter, maximizing both sensitivity and specificity. A GSVA score cutoff of - 0.195 was utilized to define “bivalent-low” and “bivalent-high” tumors across all three included data sets. The Kaplan-Meier method was used to determine EFS and OS. Survival analyses were performed in R using the “survival” package (version 3.5-3) ^67^. Differences in survival between groups was assessed with the log-rank test.

To determine the association between bivalent gene expression and OS, univariate and multivariable Cox proportional hazards regression models were utilized. The proportional hazards assumption was validated for all included models through an evaluation of Schoenfeld residuals. Stratification of Cox models was implemented for variables that violated the proportional hazards assumption. All included hypothesis tests were two-sided, and p-values lower than 0.05 were considered statistically significant. Included data were assumed to be normally distributed; however, this assumption was not formally validated.

## AUTHOR CONTRIBUTIONS

MC and MA designed the study and wrote the manuscript. KM, RT, TW, RL performed experiments. MC and SV retrieved sequencing data and performed bioinformatics and statistical analyses. MC, GC, AC, CH, AP, MA assisted with data interpretation. MA provided clinical insights on the project. MC, KM, GC, SV, RT, TW, RL, AC, CH, AP, MA reviewed, edited, and approved the submitted manuscripts.

## ACKNOWLEDGEMENTS

This work was supported in part by the Burroughs Wellcome Fund Early Scientific Training Program to Prepare for Research Excellence Post-Graduation (BEST-PREP; MC), University of Chicago Pritzker School of Medicine Summer Research Program (SRP; MC), a kind gift from Barry & Kimberly Fields (CH). Also, supported by the National Institutes of Health R37CA262781 (MAA), the Alex’s Lemonade Stand Foundation (MAA), and the Ludwig Center at the University of Chicago (CH). CH is a Howard Hughes Medical Institute Investigator. The contents are solely the responsibility of the authors and do not necessarily represent the official views of the NIH.

## COMPETING INTERESTS

CH is a shareholder of Shanghai Epican Genetech Co. Ltd. that licensed 5-hmC-Seal from the University of Chicago. CH is a scientific founder and scientific advisory board member of Accent Therapeutics, Inc. MAA has performed consulting for Innervate Radiopharmaceuticals.

## DATA AVAILABILITY

ChIP-Seq, 5-hmC, KAS-Seq, and RNA-Seq raw and processed data for included neuroblastoma cell lines are available at GSE212882.

## CODE AVAILABILITY

The underlying code for this study is not publicly available but may be made available on reasonable request from the corresponding author.

## Notes

### Summary of Updates

This version of the manuscript has been revised to ensure high quality images are available. No changes to the context of the text or figures have been made.

