## Supplemental Figures for "5-hydroxymethylcytosine profiling of cell-free DNA identifies bivalent genes that are prognostic of survival in high-risk neuroblastoma"

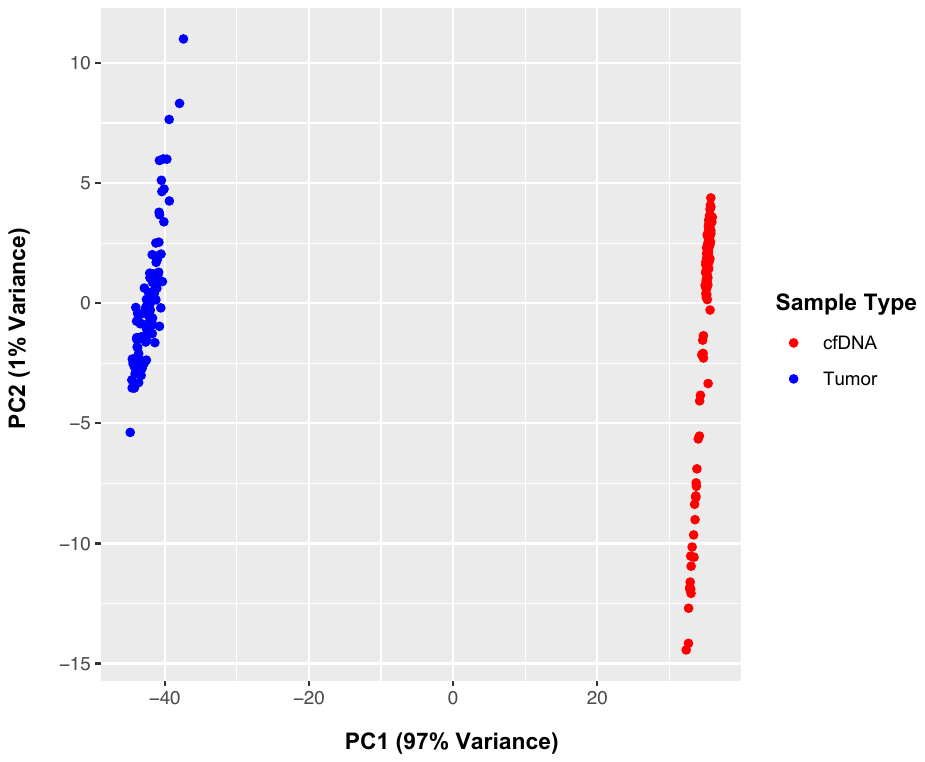
**SUPPLEMENTARY FIGURES**

**D**

**B**

**A**

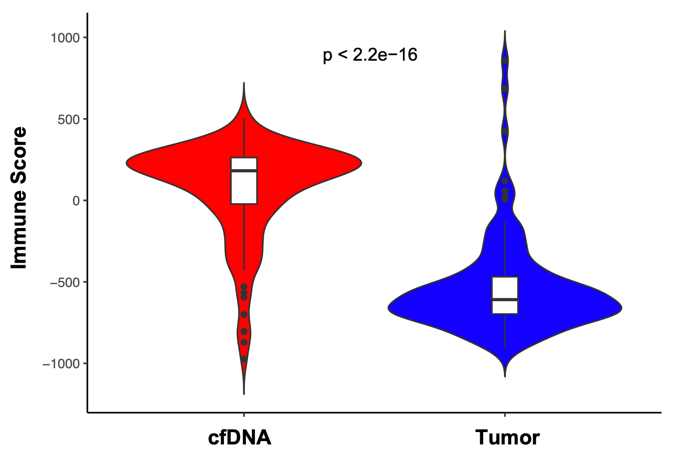

**C**

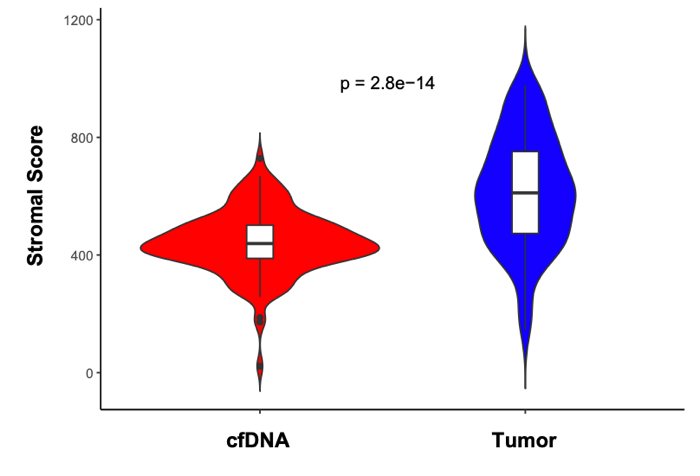

**SUPPLEMENTAL FIGURE 1: Characterization of the cfDNA and tumor biopsy derived 5-hmC profiles by Principal Component Analysis, ESTIMATE scoring, and unsupervised hierarchical clustering. A)** Principal Component analysis of cfDNA and tumor derived 5-hmC profiles shown with Principal Component 1 (x-axis) and Principal Component 2 (y-axis). **B)** Immune score and **C)** Stromal score as determined by the ESTIMATE algorithm of cfDNA and tumor 5-hmC profiles. **D)** Unsupervised hierarchical clustering of cfDNA derived 5-hmC profiles by the top 10% most significantly differentially hydroxymethylated genes identified in a comparison of cfDNA and tumor 5-hmC pofiles.

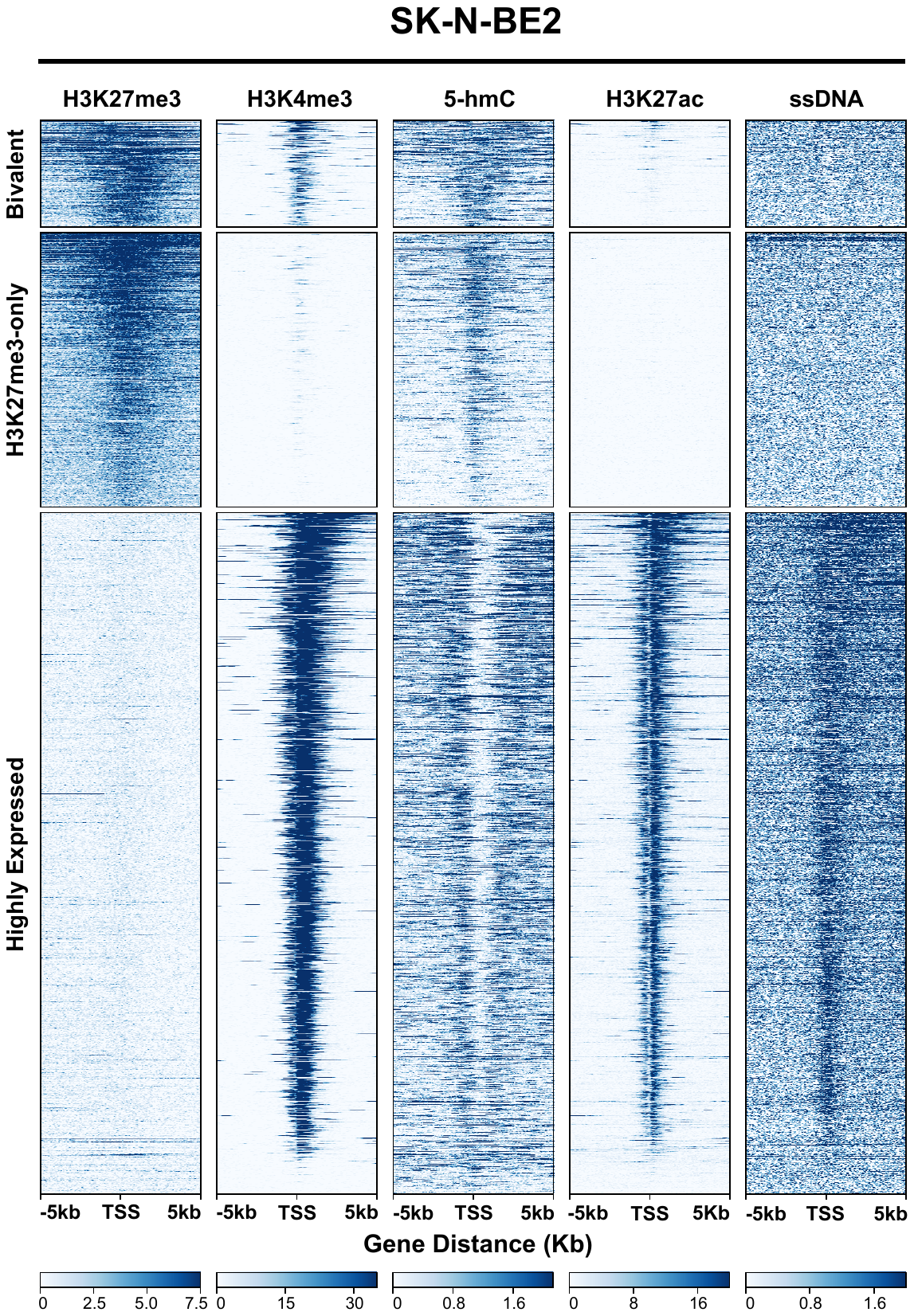

**SUPPLEMENTAL FIGURE 2: Bivalent genes are enriched for 5-hmC at the TSS in SK-N-BE2.** H3K27me3 profiles were normalized to RPKM and were visualized +/- 5kb of the TSS. Genes are ranked in descending order by relative H3K27me3 as defined in the SK-N-BE2 cell line. H3K27me3, H3K4me3, 5-hmC, H3K27ac, and ssDNA profiles (normalized to RPKM) +/- 5kb of the TSS, in the NBL-W-N cell line. Profiles were obtained across three different gene lists: bivalent (313 genes) and H3K27me3-only genes (806 genes), as defined in the SK-N-BE2 and NBLW-N cell lines, in addition to highly expressed genes (2000 genes), defined as the 2000 genes with the highest TPM in the SK-N-BE2 cell line.

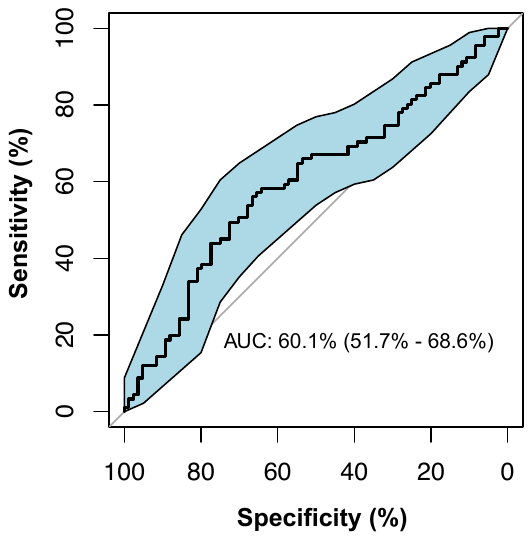

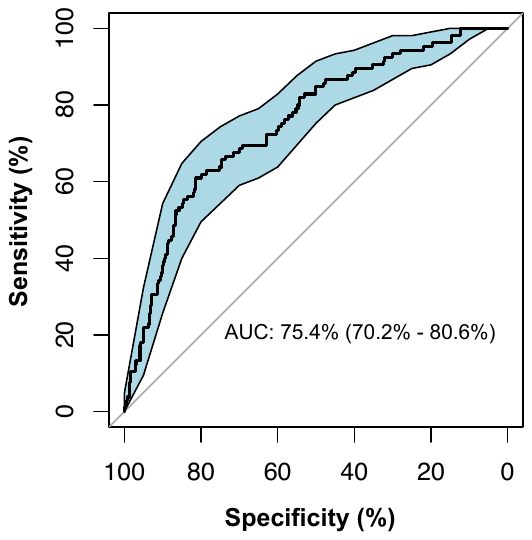

**B**

**A**

**C**

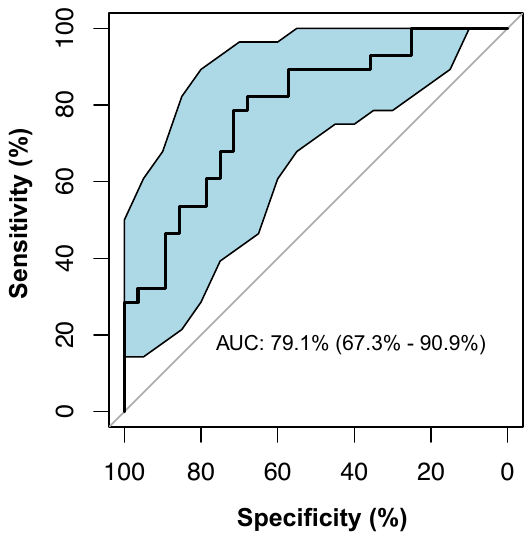

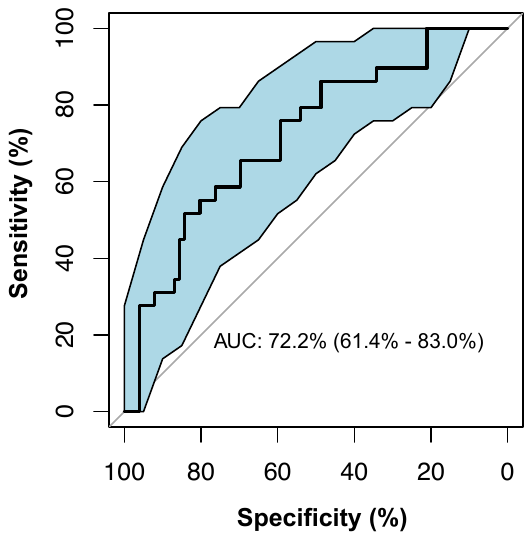

**D**

**SUPPLEMENTAL FIGURE 3: Receiver operator curves (ROC) for bivalent gene signatures against overall survival (OS).** ROC for bivalent gene signatures and OS in **A)** all 493 tumors and **B)** 175 HR tumors with known *MYCN* amplification status in the SEQC-NB dataset. ROC for bivalent gene signatures and OS in all **C)** 105 tumors and **D)** 56 HR tumors in the GSE73517 dataset. AUC indicates area under the curve. 95% confidence intervals are reported in parentheses.

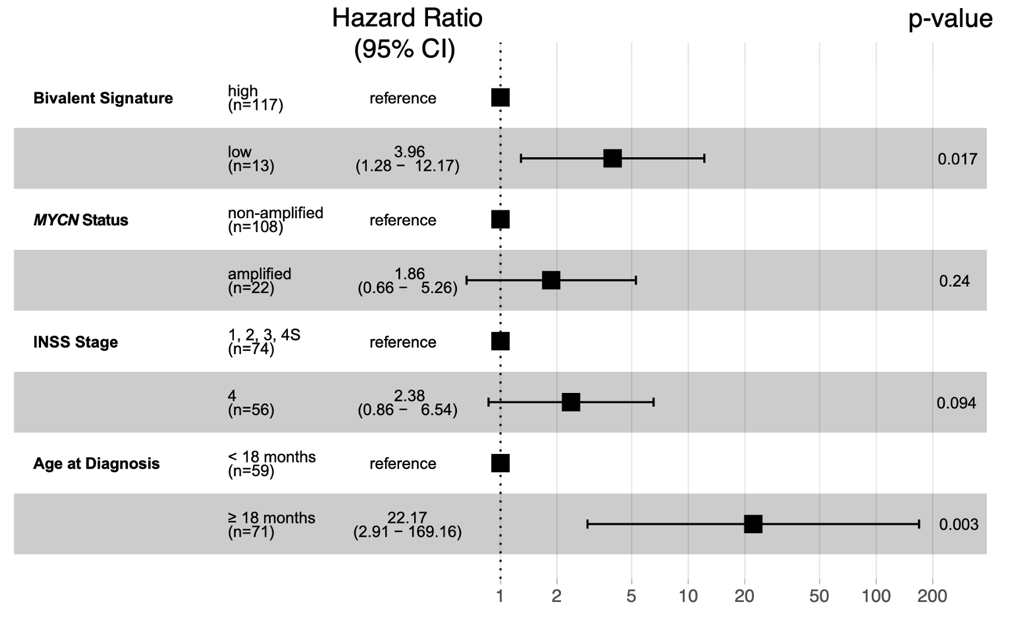

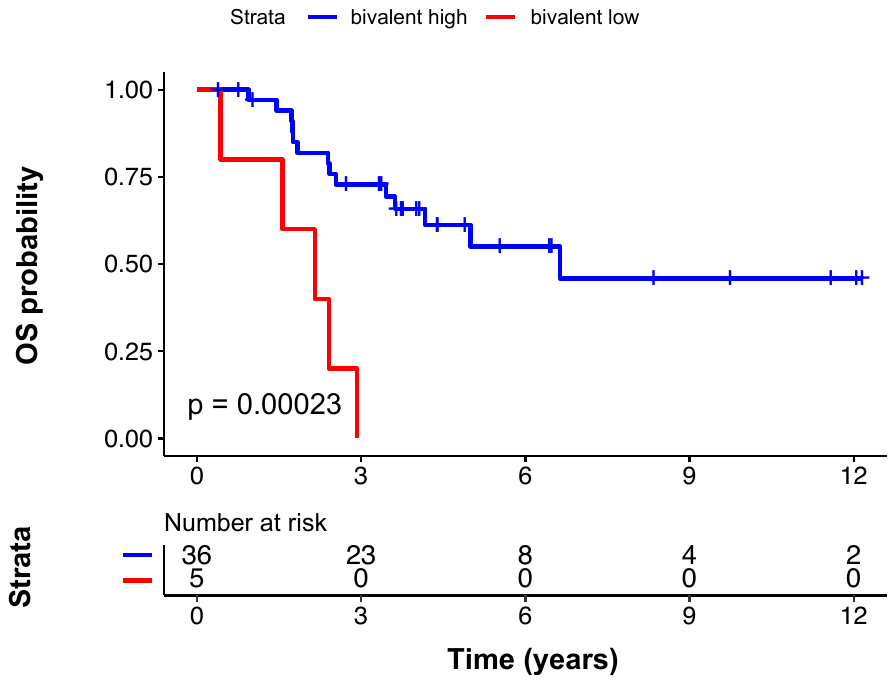

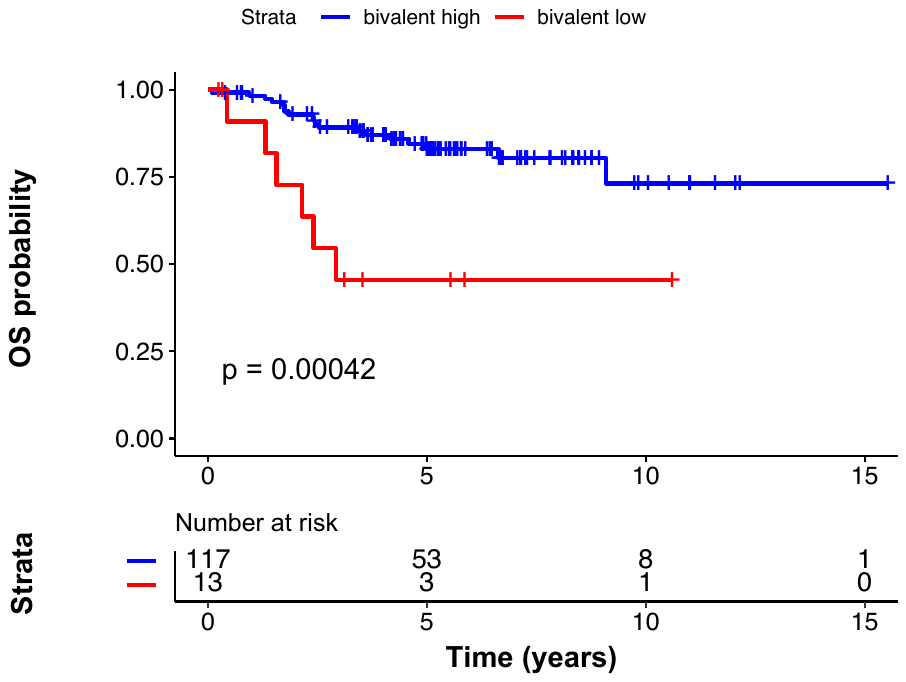

**C**

**B**

**A**

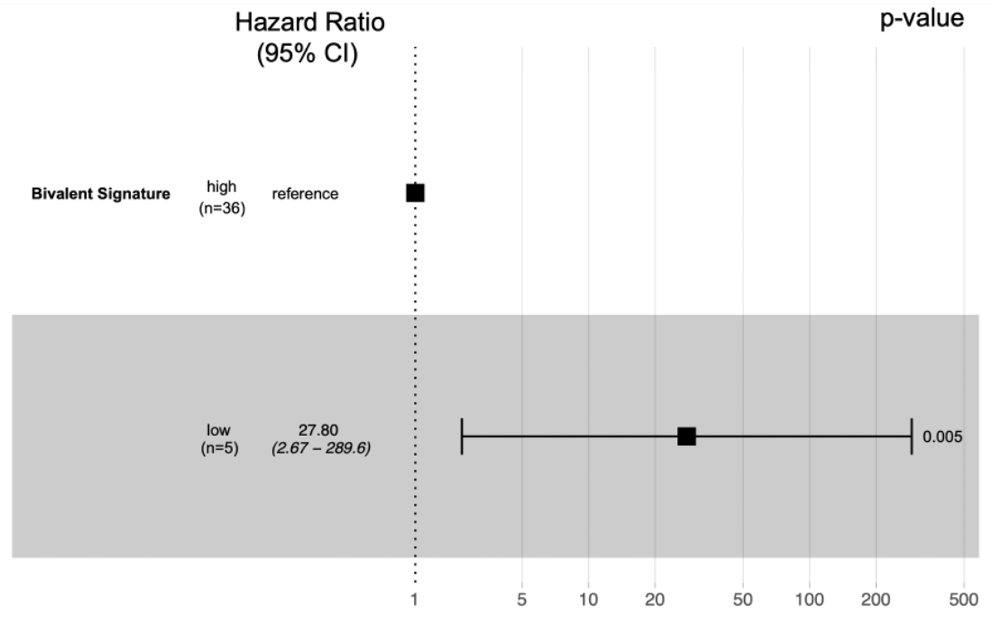

**D**

**SUPPLEMENTAL FIGURE 4: Low expression of bivalent genes is associated with poor clinical outcome in neuroblastoma patients in the E-MTAB-8248 dataset.** Expression and phenotypic data for 130 neuroblastoma patients (E-MTAB-8248) were obtained from R2. Kaplan-Meier curves depicting overall survival (OS) in all **A)** 130 patients and **B)** 41 HR patients by relative bivalent gene expression. **C)** Multivariable Cox regression analysis for OS in all 130 patients. **D)** Multivariable Cox regression analysis for OS in 41 HR patients stratified by *MYCN*-amplification status.
